# Hormone-induced thrombosis is mediated through non-canonical fibrin(ogen) aggregation and a novel estrogen target in zebrafish

**DOI:** 10.1101/2024.11.13.623199

**Authors:** Xinge Yu, Murat Yaman, Queena Y. Zhao, Sylvia M. Emly, Jacqueline K. Lee, Hongyu Su, Allison C. Ferguson, Chandrasekaran Nagaswami, Saireudee Chaturantabut, Wolfram Goessling, John W. Weisel, Richard J. Auchus, Jordan A. Shavit

## Abstract

Venous thrombosis is a well-known complication of estrogen exposure, with nearly every woman at risk across her lifetime through contraception, pregnancy, or hormone therapy. Although estrogens alter expression of coagulation factors, the mechanisms that mediate estrogen-induced thrombosis are poorly understood, partially due to the absence of an animal model. Identification of these mediators is central to understanding of hormone-induced pathophysiology, could ascertain patients at higher risk for thrombosis, and pinpoint future therapeutic targets. The zebrafish is characterized by external development, high fecundity, optical transparency, and hemostasis is highly conserved with humans. Through a transgenic line that generates GFP-tagged fibrinogen, we show rapid onset of thrombosis after exposure to various estrogens, but not progestins or testosterone. Thrombi are localized to the venous system with evidence for clot contraction. Thrombosis is only partially impeded by anticoagulants, occurs in the absence of factor VII, factor X, and prothrombin, but is dependent on tissue factor and fibrin(ogen). Finally, targeting of all known estrogen receptors does not eliminate thrombosis. The inability to completely inhibit thrombosis through genetic/pharmacologic anticoagulation or estrogen receptor disruption suggests mechanisms different from canonical coagulation/thrombosis. These studies suggest that estrogen-induced thrombosis is a unique entity distinct from other forms of venous thrombosis.

## Introduction

Venous thromboembolism (VTE) is caused by the formation of pathologic blood clots in major veins, resulting in deep vein thrombosis and/or pulmonary embolism. According to the Centers for Disease Control, VTE affects ∼900,000 people/year in the U.S., and accounts for 60,000-100,000 deaths (1). Major causes are inherited or acquired hypercoagulable states and estrogen exposures, the latter most prominently from combined oral contraceptives (COCs) (1–3), but also including pregnancy and hormone replacement therapies. Estrogen is a longstanding well-known risk factor for VTE (4–6), especially in combination with other acquired risk factors, such as smoking and obesity (7). Despite multiple clinical studies, the mechanisms of estrogen-induced thrombosis are minimally understood (8).

Estrogens target multiple receptors with various agonistic affinities for both classic nuclear hormone receptors (NHRs, the estrogen receptors (ESRs)) (8) and a G-protein coupled estrogen receptor (GPER1) (9). In premenopausal women, 17α-ethinylestradiol (EE2), a synthetic estrogen with four to five times the potency of natural estrogens, is the one most commonly used in COCs. Activities are also dependent on the crosstalk between various hormone receptors, including progestins, androgens, and estrogens (10). For example, some androgens, such as 19-nortestosterone, have weak estrogenic activities (11), and some progestins are known to have weak androgenic activities (12). Estrogen functions through two mechanisms, genomic and non-genomic (13). ER_α_ and ER_β_ (genes *ESR1* and *ESR2*) are the two NHRs, with zebrafish orthologs Esr1, Esr2a, and Esr2b (14, 15). After interacting with estrogen, these NHRs dimerize and translocate to the nucleus to bind estrogen response elements (EREs) and regulate gene transcription (16). Non-genomic effects are mediated by GPER1 (17). There is also evidence for additional receptors (18, 19), as well as new functions for known NHRs (20, 21).

Over the years, studies have shown that estrogens are able to upregulate gene expression of multiple blood coagulation factors in humans, mice, and zebrafish (8, 22–24) and to downregulate expression of anticoagulant factors (25, 26). Recent progress has been made through *in vitro* and *ex vivo* studies. In endothelial cells, 17β-estradiol (E2) activates exocytosis through ER_α_ (27), but COCs do not affect endothelial procoagulant activity, even with overexpression of the *ESR* genes (28). *Ex vivo*, E2 appears to affect clot dynamics, including formation, propagation, strength, structure, and fibrinolysis. However, *in vivo* murine studies have been largely unsuccessful, as EE2 increases thrombin generation, but there is no increase in tissue fibrin deposition or a visible thrombotic phenotype (29). Additionally, these studies show suppression of thrombosis by EE2 (30). Thus, absence of a relevant animal model has contributed to a continued lack of mechanistic understanding.

As a vertebrate model, zebrafish (*Danio rerio*) exhibits unique characteristics such as external fertilization, rapid development, high fecundity, and optical transparency, and possesses all the key coagulation factors found in humans (31–33). The transparency of zebrafish larvae allows for real-time observation of blood circulation and fluid dynamics. Our previous research utilizing genome editing in zebrafish has revealed remarkable similarities between the pro- and anticoagulant factors in zebrafish and those in humans (34–39). Zebrafish also demonstrate the preservation of major estrogen receptors and estrogen-regulated gene activities (40–42). In this study, we show that zebrafish develop venous thrombosis in response to multiple estrogens, but not progestins or testosterone. These thrombi form even in the absence or inhibition of the coagulation cascade, as well as absence of all known estrogen receptors, although tissue factor (TF, *f3* gene) and fibrin(ogen) are required. These data suggest the presence of alternative pathway(s) for thrombosis, which might involve fibrinogen crosslinking.

## Results

### E2 causes thrombosis in vivo

We first tested endogenous estrogen activity by incubating E2 for 5 hours in *fgb-egfp* larvae, and discovered fluorescent thrombi appearing prominently in the posterior cardinal vein (PCV) (Figure 1A). A semi-quantitative grading system (Figure 1B) was developed and found to be highly reproducible between independent observers. When we compared these results to quantitative measurement of fluorescence through corrected total cell fluorescence (CTCF), there was strong correlation between the two methods (Spearman’s ρ: 0.84, Figure 1C). Given this result and ease of use, all further data were collected using the grading system. The most efficacious results were observed at relatively high concentrations of E2 (50-75 µM). Subsequently, we measured larval plasma concentrations and found that only a small fraction of E2 was absorbed in individual larvae (Figure 1D).

**Figure 1:**
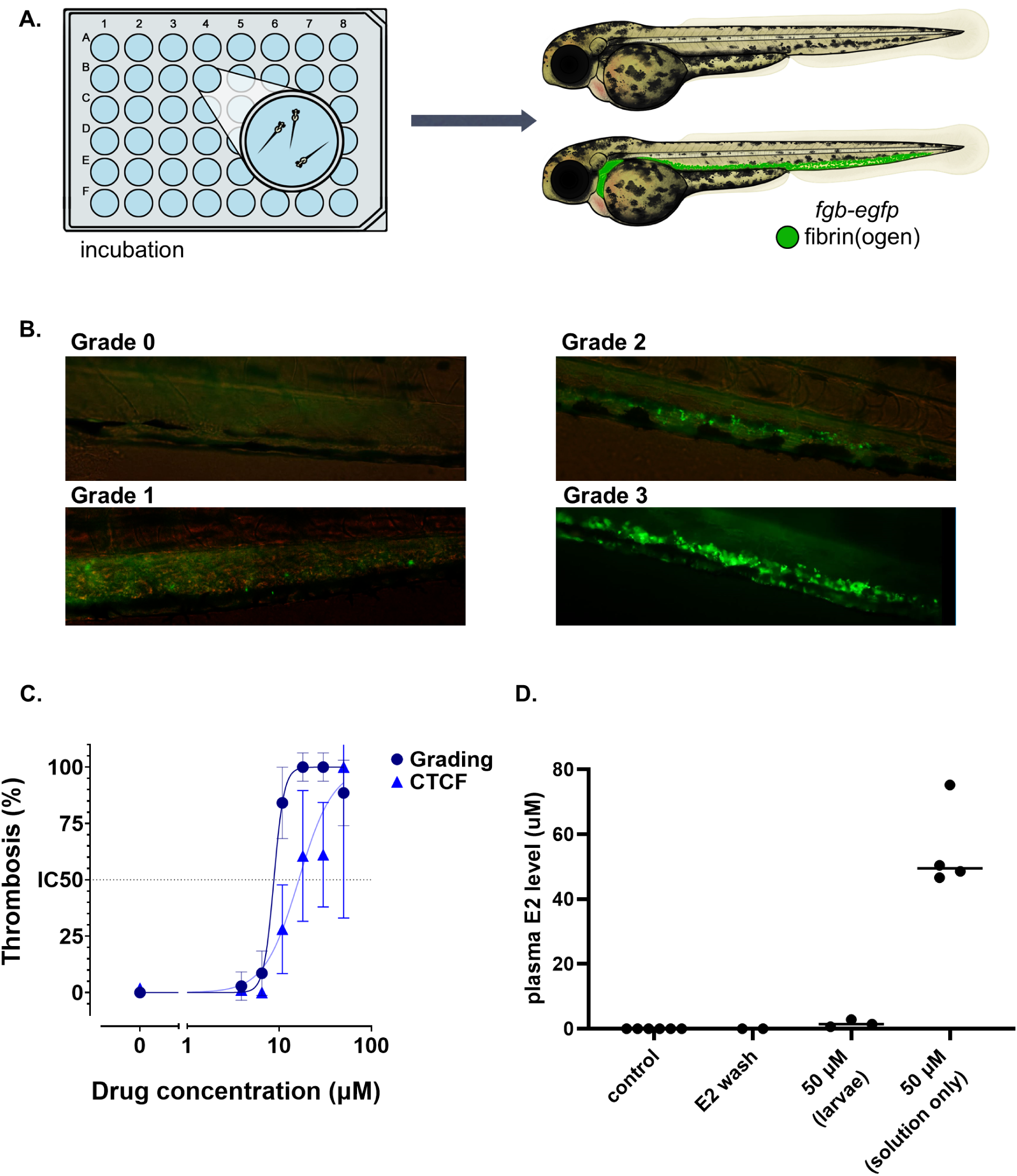
Induction and quantification of thrombosis in zebrafish larvae. (A) Estrogenic compounds were dissolved in DMSO and added to multiwell plates with E3 embryo buffer. *fgb-egfp* transgenic larvae, which generate GFP-tagged fibrin(ogen), were added at 3-5 dpf and incubated up to 48 hours. After incubation, larvae were briefly washed with system water and then mounted in 0.8% agarose and evaluated with a compound microscope and 20x objective. After incubation, estrogen-treated fish developed GFP-labeled thrombosis in the PCV, indicating fibrin(ogen) deposition, which was graded. (B) Representative images for the semi-quantitative grading system. Scoring was always performed by an observer blinded to condition and/or genotype as follows: Grade 0: no fluorescence accumulation. Grade 1: small sporadic fluorescent puncta seen scattered in the PCV. Grade 2: continuous fluorescence spreading across the PCV. Grade 3: strong fluorescence visualized across the PCV. (C) Quantification of fluorescence by CTCF in comparison with semi-quantitative grading. Each point represents the mean values from n=9-12 larvae, with error bars showing 95% confidence intervals. (D) Quantification of plasma levels of 17β-estradiol (E2) after treatment in 25 and 50 µM E2 and controls. Larvae were bled into buffer and then pooled for mass spectrometry. “E2 wash” indicates larvae washed with 50 µM E2 and then system water, without prior incubation in E2. “50 µM (solution only)” is quantification of the E2 containing buffer without larvae. Each point represents a pool of 10 larvae.

### Synthetic estrogens induce in vivo thrombosis

E2 and EE2 (the most common form of estrogen in COCs) both showed dose-dependent induction of thromboses in our model (Figure 2, A and B); however, these findings were inconsistent between experiments. What is shown are the “representative” best data, but often times we found the overall frequency of thrombosis to be less than 50%, even at the high concentrations. Therefore, we evaluated two prodrugs that are metabolized to EE2 *in vivo*, mestranol and quinestrol. These compounds demonstrated greater potency and efficacy, as they were consistent across multiple experiments, although required longer incubations (24-48 hours, Figure 2, C and D).

**Figure 2:**
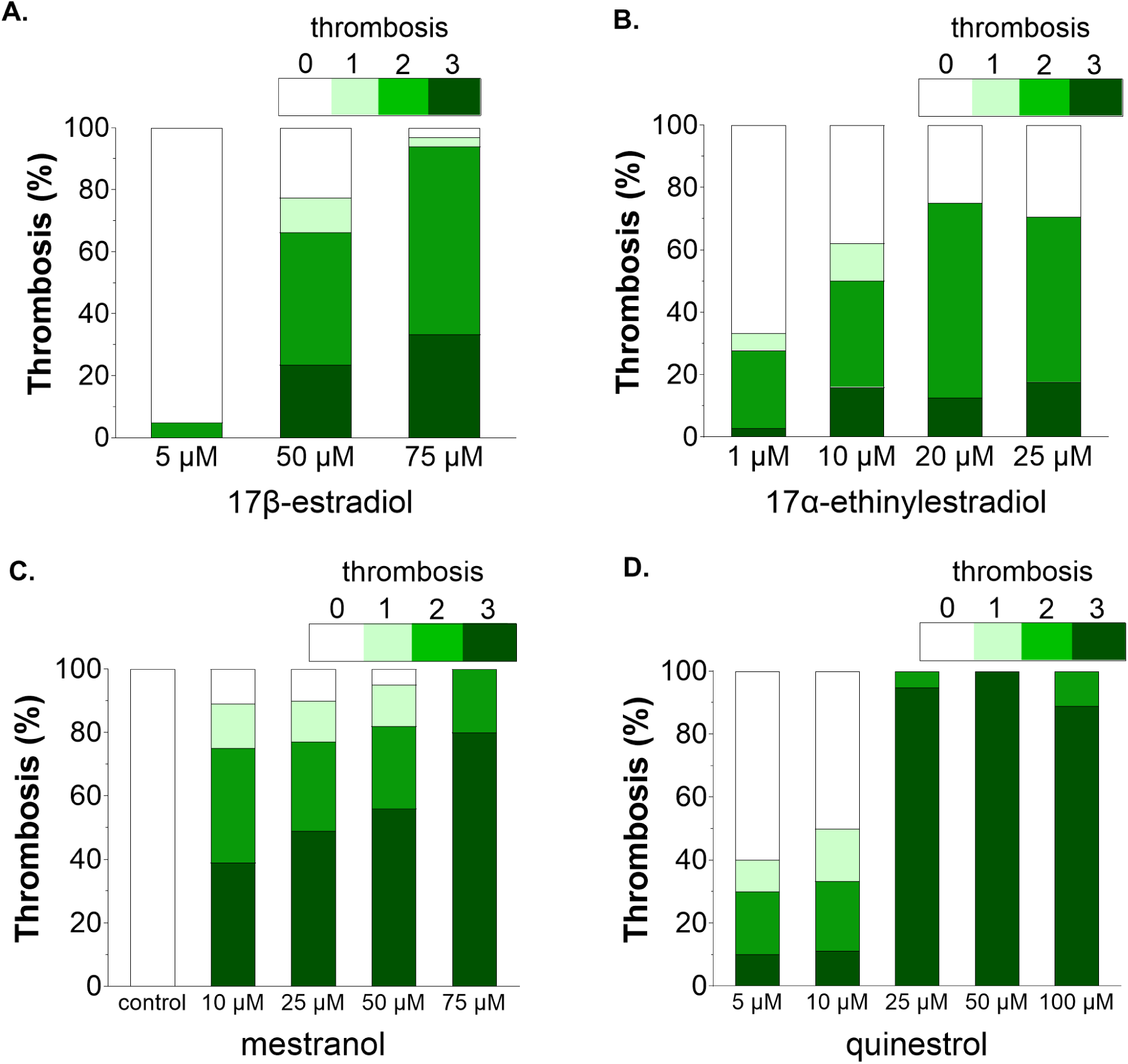
Estrogen analogs induce dose-dependent *in vivo* thrombosis. Dose-dependent increases in thrombosis were observed with multiple estrogens. (A) 17β-estradiol (E2) treatment for 5 hours at 5 dpf. n=41, 234, 33. (B) 17α-ethinylestradiol (EE2) treatment for 24 hours at 4 dpf. n=36, 50, 8, 17. (C) Mestranol treatment for 24 hours at 4 dpf. n=24, 38, 39, 39, 20. (D) Quinestrol treatment for 24 hours at 4 dpf. n=10-19 in each group. All incubations were performed in the *fgb-egfp* transgenic background and data collected by an observer blinded to condition.

### Testosterone derivatives and progestins do not cause thrombosis

Testosterone has been shown to have an association with increased VTE (43), so we evaluated three testosterone derivatives. None of these showed any significant thrombotic activity (Figure 3, A-C). Progestins are considered to be relatively safe when taken independently (8), and four progestins (at the maximum tolerated doses) across different generations were evaluated, but none of them triggered thrombosis (Figure 3D).

**Figure 3:**
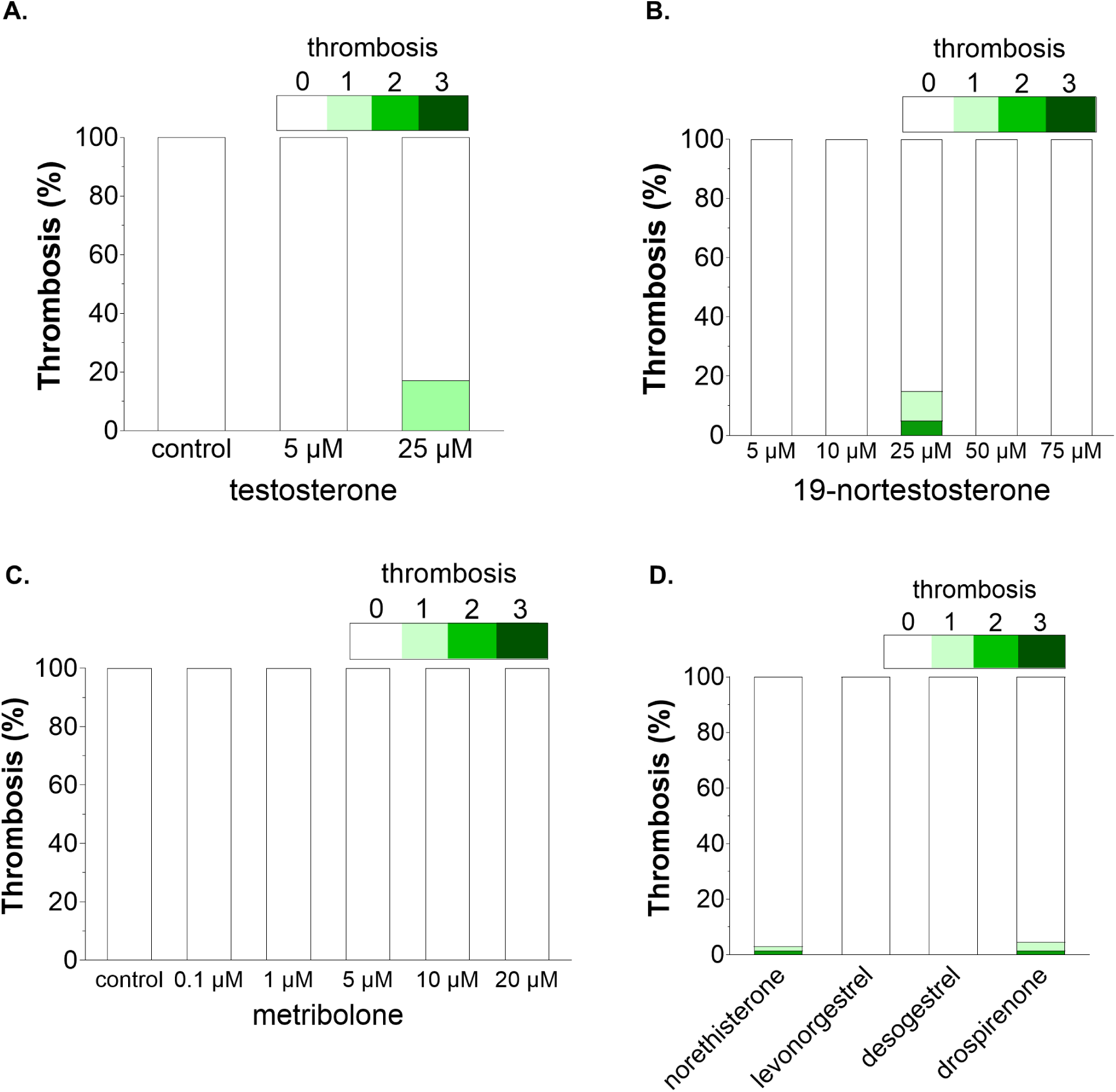
Testosterone derivatives and progestins do not cause thrombosis. (A) Testosterone incubations were performed for 24 hours starting at 4 dpf. (B) 19-nortestosterone incubations were for 48 hours starting at 3 dpf. (C) Metribolone incubations were 24 hours starting at 4 dpf. n=10-30 for each testosterone derivative concentration. (D) Multiple progestins were evaluated at the maximum tolerated dose for 5 hours on 5 dpf. Norethisterone 10 µM, levonogestrel 25 µM, desogestrel 1 µM, drospirenone 25 µM. n=30-65 for each group. All data were collected in the *fgb-egfp* transgenic background by an observer blinded to condition.

### Static and time-lapse imaging of estrogen-induced thrombosis

To confirm the localization and composition of estrogen-induced thrombi, we used confocal and scanning electron microscopy. Confocal imaging was performed in a double transgenic that included *flk-mcherry*, labeling endothelial cells with red fluorescence, along with *fgb-egfp* to identify thrombi. GFP was not colocalized with endothelial cells, indicating its presence in the lumen of the PCV (Figure 4, A and B, and Supplemental Video). Additionally, GFP was not observed in the dorsal aorta (DA), although was occasionally found in the intersegment vessels (ISVs). These details indicate that thromboses were not generated within nor internalized by endothelial cells. Increasing doses or duration of incubation did not result in arterial thrombosis.

In humans, it is well-known that thrombi often originate at a site of injury, and then extend through the vasculature. However, given the spontaneous and unpredictable nature of estrogen-induced thrombosis, it is unknown how this occurs in patients. Time-lapse imaging was performed over a six-hour period after initiation of treatment with E2. The results show that thrombosis occurs simultaneously across the venous endothelium, rather than initiating and spreading from a single location (Figure 4, C-F). Scanning electron microscope images revealed the morphology of these thrombi in E2-treated larvae (Figure 4, G and H). We observed polyhedrocytes (Figure 4H), consistent with the shape change that occurs in erythrocytes after clot contraction(44, 45). We have previously shown that this process occurs in zebrafish thrombi that develop after induced endothelial injury (46). Sparse fibers were also observed (Figure 4H), which we hypothesize could be composed of fibrin and/or fibrinogen. Given these findings, we evaluated cellular effects of erythrocytes and thrombocytes, and assessed the influence of von Willebrand factor (*vwf*). The *mon^tg234^* larvae, which lack erythrocytes, surprisingly showed a slightly increased incidence of thrombosis after estrogen exposure, suggesting that red cells potentially buffer against estrogen-induced thrombus formation (Figure 4I). Loss of VWF also did not affect estrogen-induced thrombosis, as *vwf-*null larvae formed wild-type levels of thrombosis (Figure 4J). Using the *cd41-egfp* thrombocyte reporter line, we counted thrombocytes that became immobilized in the PCV after mestranol treatment, and there was no difference, indicating a lack of thrombocyte recruitment to the site of thrombi (Figure 4, K and L).

### Estrogen-induced thrombi are not dependent on the common pathway of coagulation

To further determine the origins of the thrombi, we evaluated coagulation using inhibitors and multiple clotting factor knockout lines. We initially tested several clinical anticoagulants, including warfarin, rivaroxaban (factor Xa inhibitor), and dabigatran (thrombin inhibitor), and knockouts of factor X (*f10*) and prothrombin (*f2*) (35, 39). We have previously shown both the inhibitors and knockouts result in complete loss of induced thrombosis secondary to endothelial injury (34, 35, 39). In contrast, we found that the anticoagulants significantly reduced the intensity and frequency of estrogen-induced thrombosis, but did not completely block its occurrence (Figure 5, A and B). Surprisingly, in the *f10* and *f2* knockouts, which would be expected to have no thrombus formation, there was no inhibition of thrombosis in the homozygous mutants (Figure 5, C and D). Given these results, we evaluated whether fibrinogen itself is a component in the observed thrombi by examining estrogen-induced thrombosis in an a-chain (*fga*) knockout. Absence of the a-chain has proven to result in complete loss of fibrinogen in mammals (47–49) and zebrafish (36, 50). We have shown that the *fga* knockout also has complete absence of endothelial injury-induced thrombosis (36). Siblings from *fga^+/-^* incrosses were incubated with either E2 or mestranol, and in both cases there was no thrombosis detected (Figure 5, E and F). The *fgb-egfp* line is labeled with GFP on the β-chain, therefore loss of fluorescent thrombi in the a-chain knockout confirms that these are fibrin(ogen)-containing structures rather than non-specific GFP accumulation.

**Figure 4:**
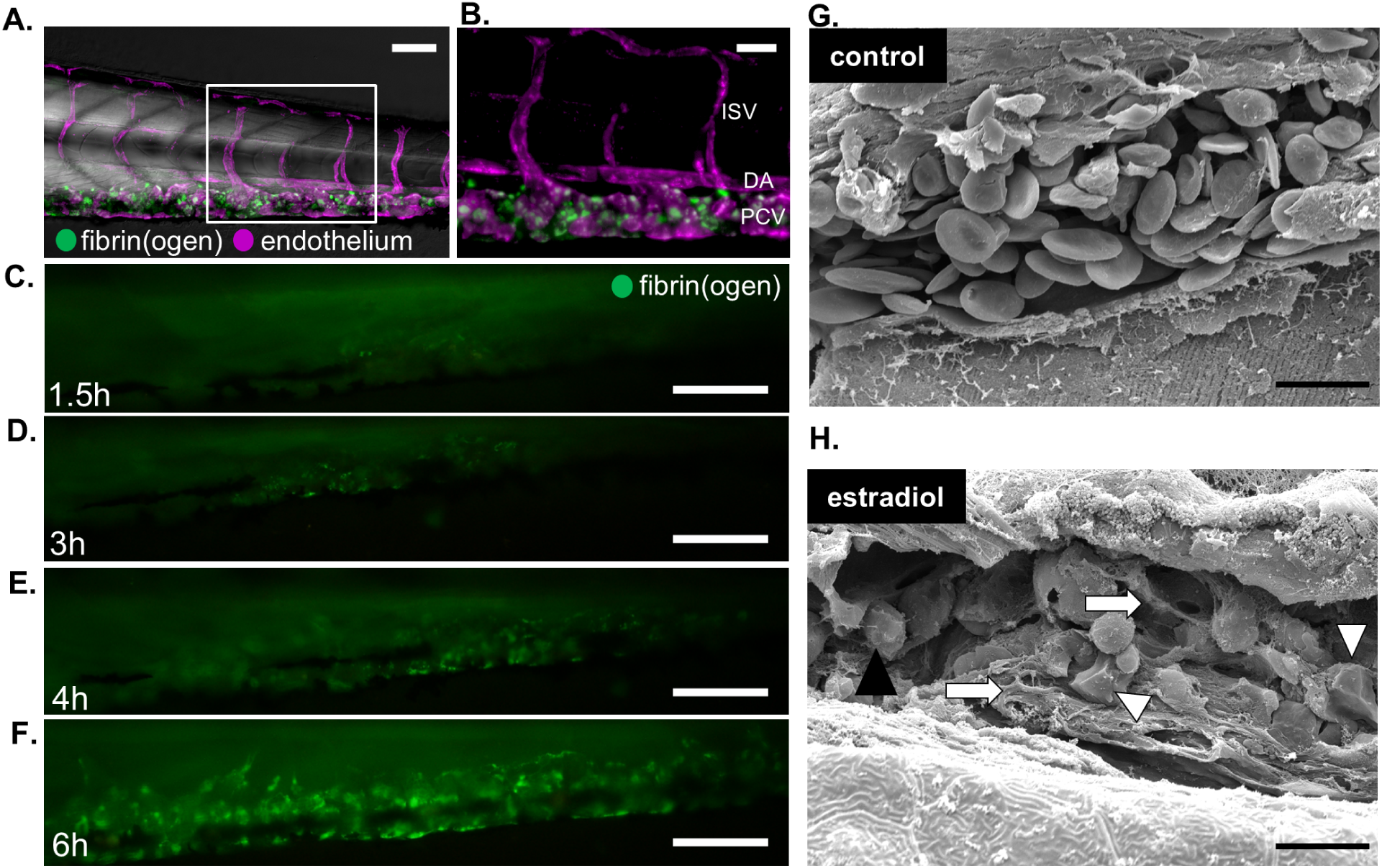
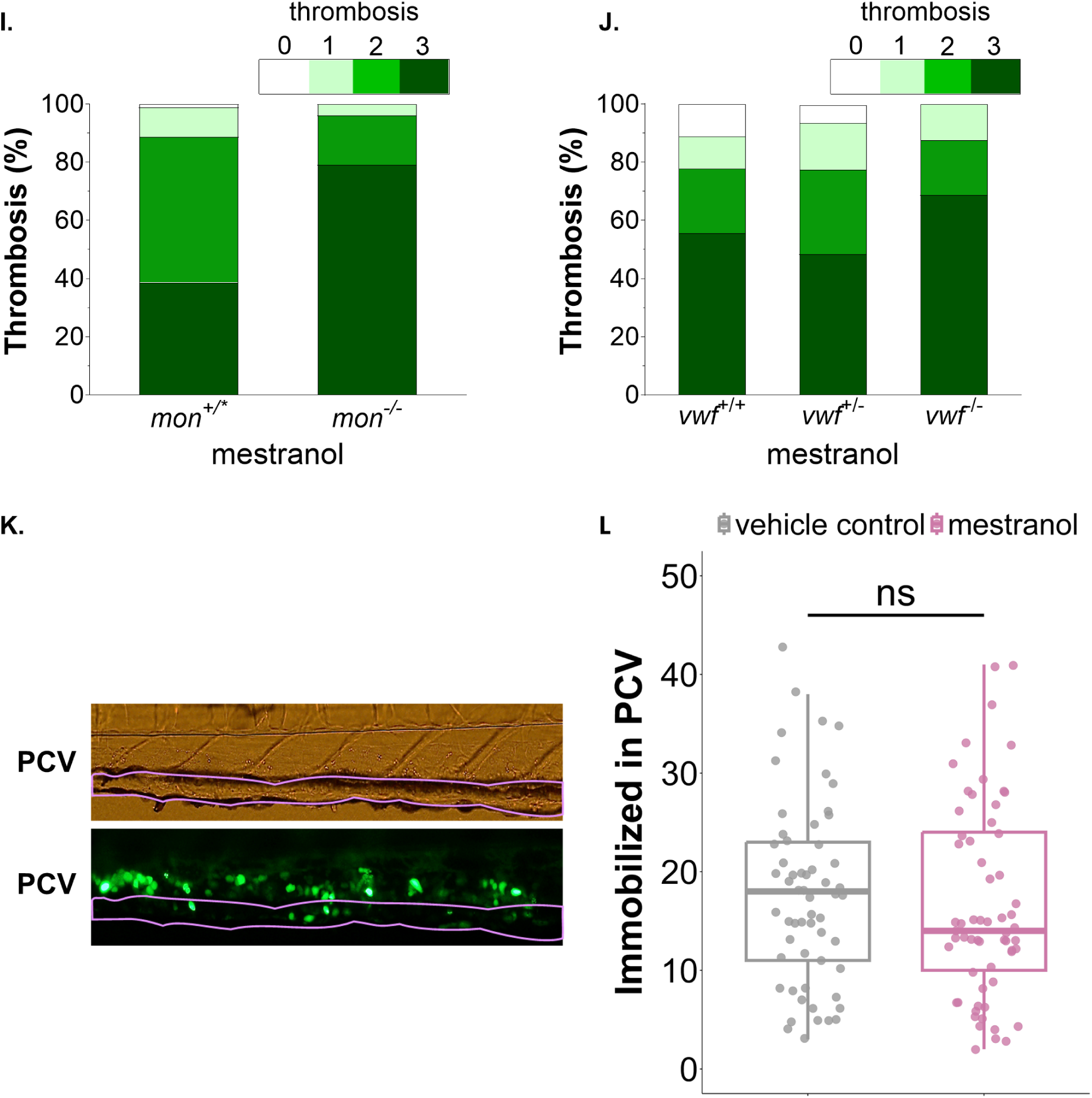
Static and time-lapse imaging of estrogen-induced thrombosis. (A-B) Fish larvae were from an *fgb-egfp*;*flk-mcherry* hybrid background, false colored to represent thrombi in green and endothelial cells in magenta. Confocal microscopy demonstrates that the GFP-labeled thrombi were contained within the PCV, and not the dorsal aorta (DA) or intersegment vessels (ISV). Scale bars, (A) 100 µm, (B) 20 µm. (C-F) Compound microscopy time-lapse images of thrombosis formation over 6 hours reveal initiation simultaneously along the PCV, rather than from a single location. Scale bars, 100 µm. (G-H) Scanning electron microscopic images comparing the PCV of a DMSO (control) treated fish (G) and a grade 2 larvae treated with E2 (H). In the control group, erythrocytes exhibit typical flat and oval morphology. In the treated larva, scattered fibers that may be fibrin(ogen) are seen attached to erythrocytes (arrows). Polyhedrocytes (arrowheads) are observed, suggestive of clot contraction. Scale bars in (G-H), 10 µm. (I-J) 48 hour mestranol treatments in offspring from *mon^+/-^* incrosses (n=88 in *mon^+/*^*, mix of wild-type and heterozygotes, 24 in *mon^-/-^*; 3◊5 dpf with 5 µM mestranol) and *vwf^+/-^*incrosses (n=18, 31, 16; 2◊4 dpf with 25 µM mestranol). p < 0.005 between *mon* groups, and no statistical significance in *vwf* using Fisher exact test. (K-L) Thrombocyte counting across larvae treated with vehicle control (DMSO) and mestranol (25 µM) for 24 hours and observed at 5 dpf (n=58, 61). (K) A thrombocyte reporter line (*cd41-egfp*) was utilized and the PCV regions (magenta outline) were selected based on brightfield and fluorescence images. (L) Immobilized thrombocytes were counted in the outlined PCV regions, and box-and-whisker plots are shown (vehicle control: box=11-23, median=18, whiskers=3-38, outlier=43; mestranol: box=10-24, median=14, whiskers=2-41, no outliers). No significant difference (ns) was detected by Student t-test. Except for K and L, all data were collected in the *fgb-egfp* transgenic background. All data were collected by an observer blinded to condition.

**Figure 5:**
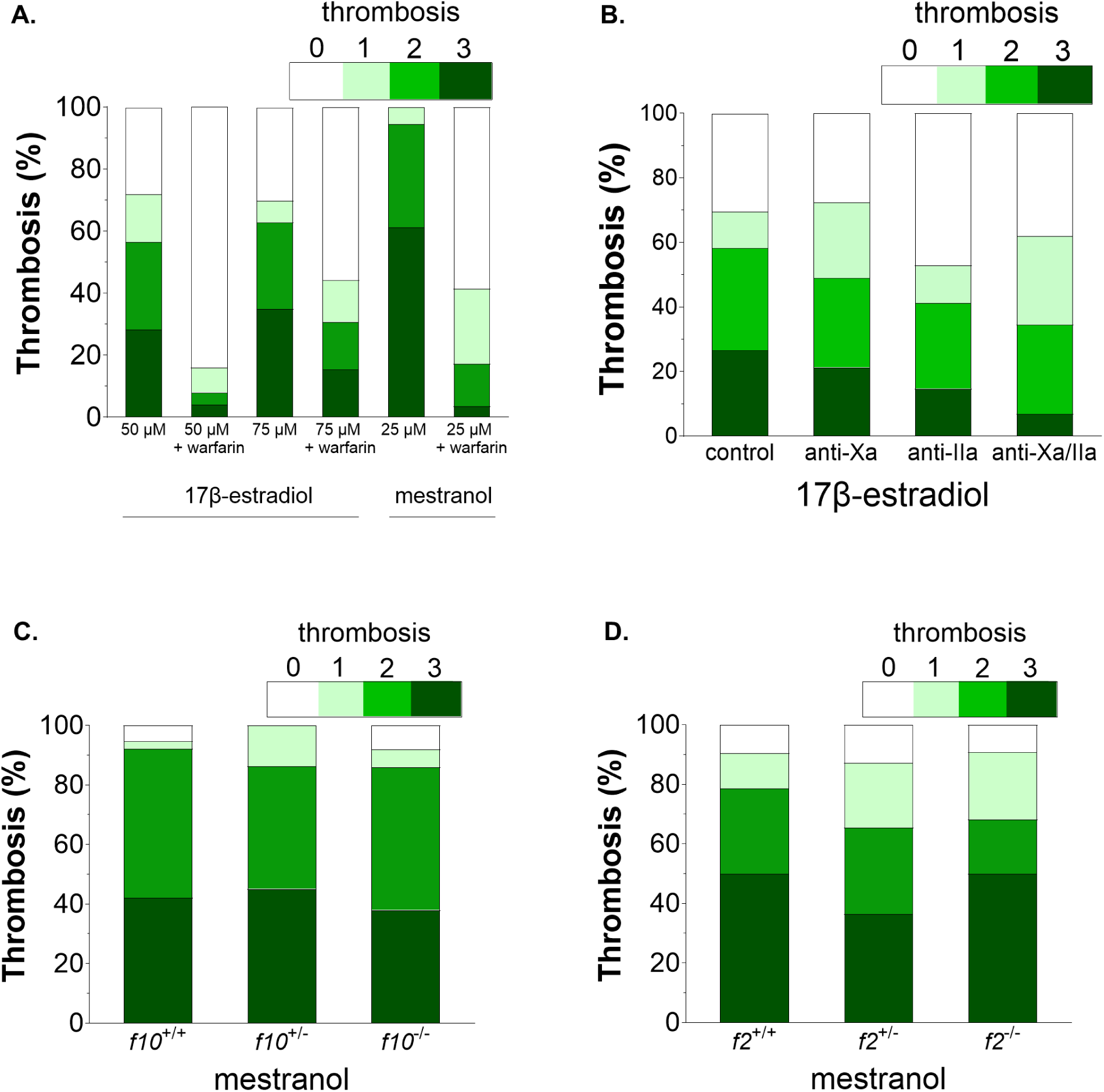
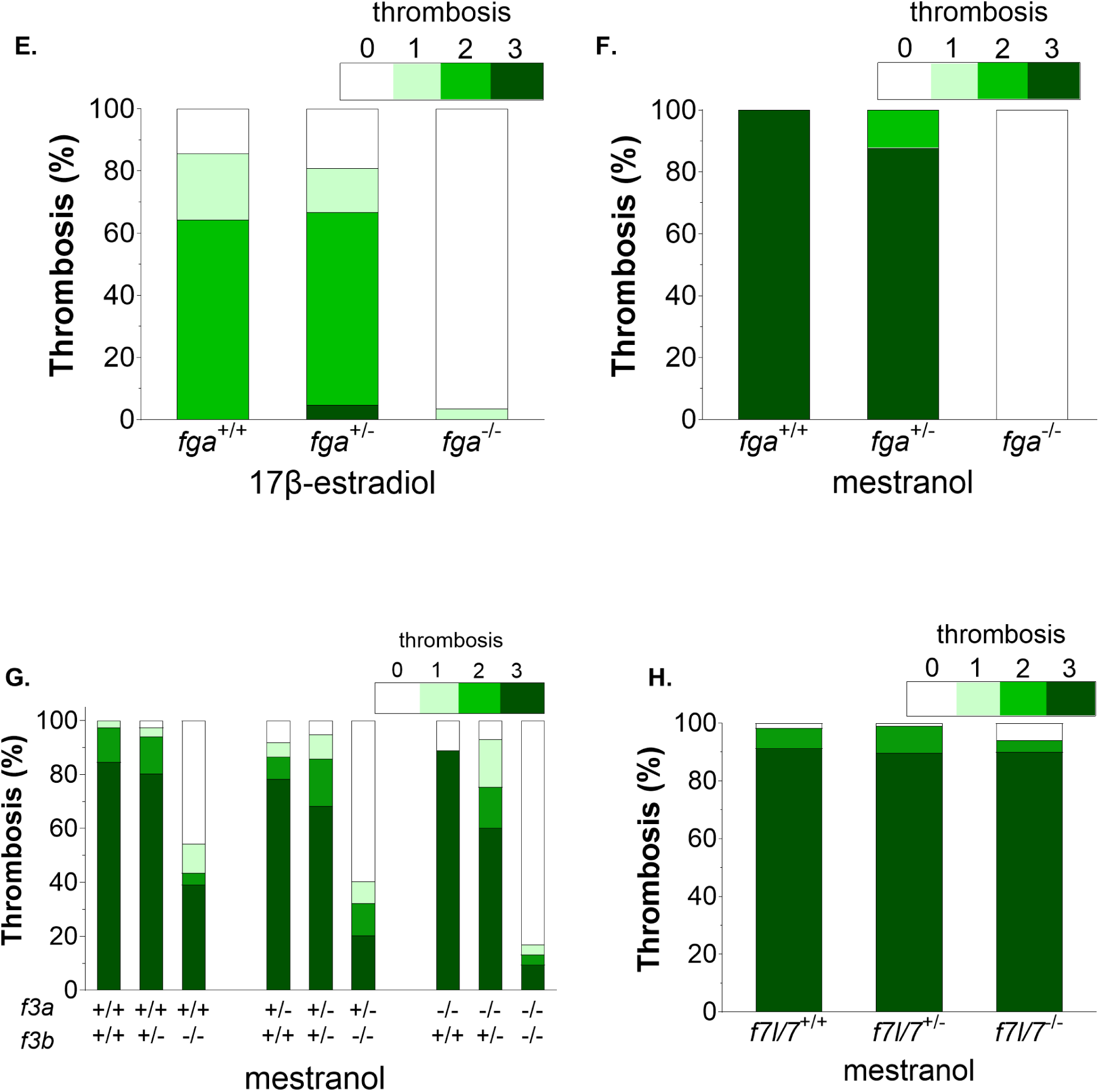
Estrogen-induced thrombi are dependent on fibrin(ogen), but not the canonical coagulation cascade. (A) Larvae were pretreated with warfarin, followed by estrogens. In the first group, warfarin (50 µg/ml) was initiated at 3 dpf, followed by E2 for 5 hours at 5 dpf. The mestranol-exposed larvae were pretreated with warfarin (10 µg/ml) at 2 dpf, followed by 25 µM mestranol for 24 hours starting at 4 dpf. n= 39, 25, 43, 52, 18, and 29. p=0.0001 for 50 µM E2 treatment, p=0.002 for 75 µM E2, p<0.0001 for mestranol. (B) Larvae were co-treated with E2 and rivaroxaban (anti-Xa) or dabigatran etexilate (anti-IIa) at 250 and 50 µM, respectively, for 5 hours at 5 dpf. n=103, 74, 50, and 51. p=0.03 (anti-Xa), 0.03 (anti-IIa), 0.001 (combination). (C-D) 48 hours of mestranol treatments in offspring from, *f10^+/-^*incrosses (n=38, 51, and 50; 3◊5 dpf with 5 µM mestranol), and *f2^+/-^* incrosses (n=42, 55, 22, 2◊4 dpf with 25 µM mestranol). No statistically significant differences were observed. (E) Offspring from *fga^+/-^* incrosses were treated with 50 µM E2 (n=14, 42, and 28 at 5 dpf for 5 hours) or (F) 25 µM mestranol (n= 18, 41, and 27 at 3 dpf for 48 hours), resulting in essentially no thrombi. p<0.0001 for both groups. (G) 25 µM mestranol treatment in *f3a^+/-^;f3b^+/-^*incrosses (n=39, 116, 92 for the *f3a*^+/+^ strata; 37, 154, 99 for the *f3a*^+/-^ strata; 27, 73, 53 for the *f3a*^- /-^ strata). p<0.0001 for each comparison of *f3b* genotypes within separate *f3a* strata, and p=0.0002 for comparison of the *f3a* genotype within the *f3b*^-/-^ genotype group, while there was no significant difference for comparison of *f3a* to the other *f3b* genotype groups. (H) Mestranol treatment in offspring from *f7l/7^+/-^* incrosses treated with 25 µM mestranol (n=58, 106, and 51; 3◊5 dpf for 48 hours) with no significant differences based on *f7l/7* genotype. All data were collected in the *fgb-egfp* transgenic background by an observer blinded to condition and/or genotype. Fisher exact test was applied for all statistical comparisons.

These data, as well as the time-lapse imaging, suggest that estrogen-induced thrombi may be composed of fibrin and/or fibrinogen, and initiated through endothelial cell signaling. TF is known for cell signaling properties, distinct from its role in coagulation (51). The gene for TF is duplicated in zebrafish, with paralogs *f3a* and *f3b* (52). Larvae devoid of *f3b* showed a ∼50% reduction in estrogen-induced thrombosis (Figure 5G). Loss of *f3a* alone had no significant effect, but double knockouts (*f3a^-/-^*; *f3b^-/-^*) exhibited an ∼80% decrease in thrombosis, demonstrating a prominent role for TF (Figure 5G). Given these data, we hypothesized that loss of factor VII (FVII) would phenocopy TF deficiency. The *f7* gene in zebrafish is triplicated (*f7*, *f7l*, *f7i*) in cis(53), and paralogs are closely related to human FVII (54–56). We and others have found that loss of FVII leads to a partial reduction in induced thrombosis due to endothelial injury (57), but combined deletion of the tandem *f7l*/*f7* locus completely ablates thrombus formation (Shavit laboratory, manuscript in preparation). *f7i* has been shown to be inhibitory rather than procoagulant and thus was not evaluated (56). Surprisingly, examination of estrogen-induced thrombosis showed no effect with loss of FVII activity (Figure 5H).

### Estrogen-induced thrombosis is mediated through a non-canonical target

Next, we explored whether the effect is mediated through known estrogen receptors. We first examined fulvestrant (58), an estrogen antagonist with similar binding affinity to E2. It is simply a 7α-alkylsulphinyl analog of E2 that prevents nuclear hormone receptor dimerization, which is essential for downstream signaling and facilitates receptor degradation. We treated zebrafish larvae with E2 or mestranol and fulvestrant, and observed significant reduction of thrombosis (Figure 6, A and B). We proceeded to examine the effect of loss of the various estrogen receptors, starting with the NHRs (*esr1*, *esr2a*, and *esr2b*). Previous studies with small indels have found that double and triple, but not single mutants have effects on reproduction (59), so we chose to produce complete locus deletions of each gene to ensure complete ablation (Supplemental Figure 1). Since *esr1* and *esr2a* are tightly linked, we had to produce these double mutants through iterative targeting. Using a triple heterozygous mutant incross, we were able to evaluate single *esr2b*, double *esr1/2a*, and triple homozygous mutants (Figure 6C), all of which developed equivalent levels of estrogen-induced thrombosis. The same was true of a G-protein coupled estrogen receptor 1 (*gper1*) knockout line (60), which also exhibited persistent thrombosis (Figure 6D). There is evidence for a different membrane-associated GPR in the mouse that responds to estrogen, which is selectively targeted by the synthetic nonsteroidal ligand STX (18, 61). STX treatment in zebrafish demonstrated no effect on thrombosis formation across multiple concentrations up to 100 µM (Figure 6E)(18). Taken together, these data suggest the existence of an alternative estrogen target that is distinct from these other known receptors.

**Figure 6:**
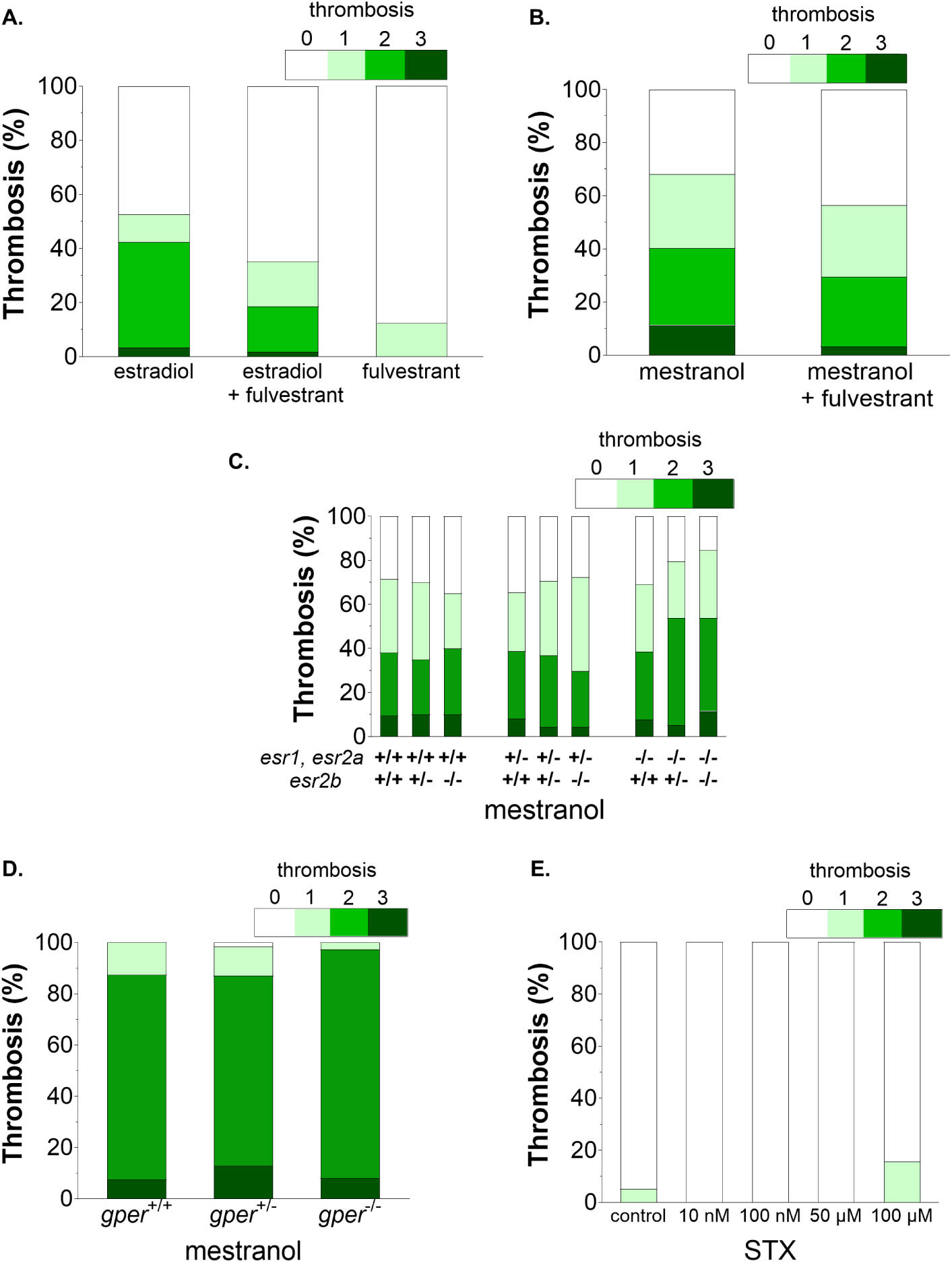
Estrogen-induced thrombosis appears to be mediated through a non-canonical estrogen target. (A) Co-treatment of E2 (50 µM) and fulvestrant (50 µM) at 5 dpf for 5 hours. n=130, 110, 8. p<0.0001 between estradiol and estradiol + fulvestrant, by Fisher exact testing. (B) Co-treatment of mestranol (2.5 µM) and fulvestrant (25 µM) at 3 dpf for 48 hours. n=69 and 156. p<0.05 by Fisher exact testing. (C) Triple heterozygous *esr1^+/-^;esr2a^+/-^;esr2b^+/-^*mutants were incrossed and larvae treated with 5 µM mestranol at 3 dpf for 48 hours (n=21, 40, 20 for the *esr1*/*esr2a*^+/+^ strata; 49, 92, 47 for the *esr1*/*esr2a*^+/-^ strata; 26, 39, 26 for the *esr1*/*esr2a*^-/-^ strata). Genotyping was performed after thrombosis quantification. Fisher exact testing revealed no significant differences. (D) Larvae from *gper*^+/-^ incrosses were treated with 5 µM mestranol at 3 dpf for 48 hours. Genotyping was performed after thrombosis quantification. No difference was detected by Fisher exact testing, n= 40, 62, and 37. (E) STX(18) treatment at 3 dpf for 48 hours. No statistical significance observed among the groups (n=19, 17, 20, 20, 20). All data were collected in the *fgb-egfp* transgenic background by an observer blinded to condition and/or genotype.

It is well known that 30-40% of zebrafish genes are duplicated due to an ancient event (62). The whole genome has been sequenced (53) and it is believed that there are only three *esr* genes, but the possibility remains that additional ones may exist. A Y549S substitution has been shown to render zebrafish Esr1 constitutively active (63). We cloned *esr1* under control of a ubiquitous promoter (64) and engineered this substitution, but transgenic embryos did not survive (data not shown). Therefore, we swapped in a heat shock promoter to enable activation of *esr1* at selected timepoints, along with a fluorescent *mcherry* reporter (Figure 7A). We injected this construct into the *ere-gfp* transgenic line, which has been shown to be responsive to activated estrogen receptors (41). After heat shock, both *mcherry* and *gfp* signals were observed indicating successful expression of the mutant *esr1* and activation of the *ere-gfp* transgene (Figure 7B). However, when injected into the *fgb-egfp* background, no thrombosis was observed in either group. Taken together, these data suggest that estrogen-induced thrombosis is mediated through a novel target.

**Figure 7:**
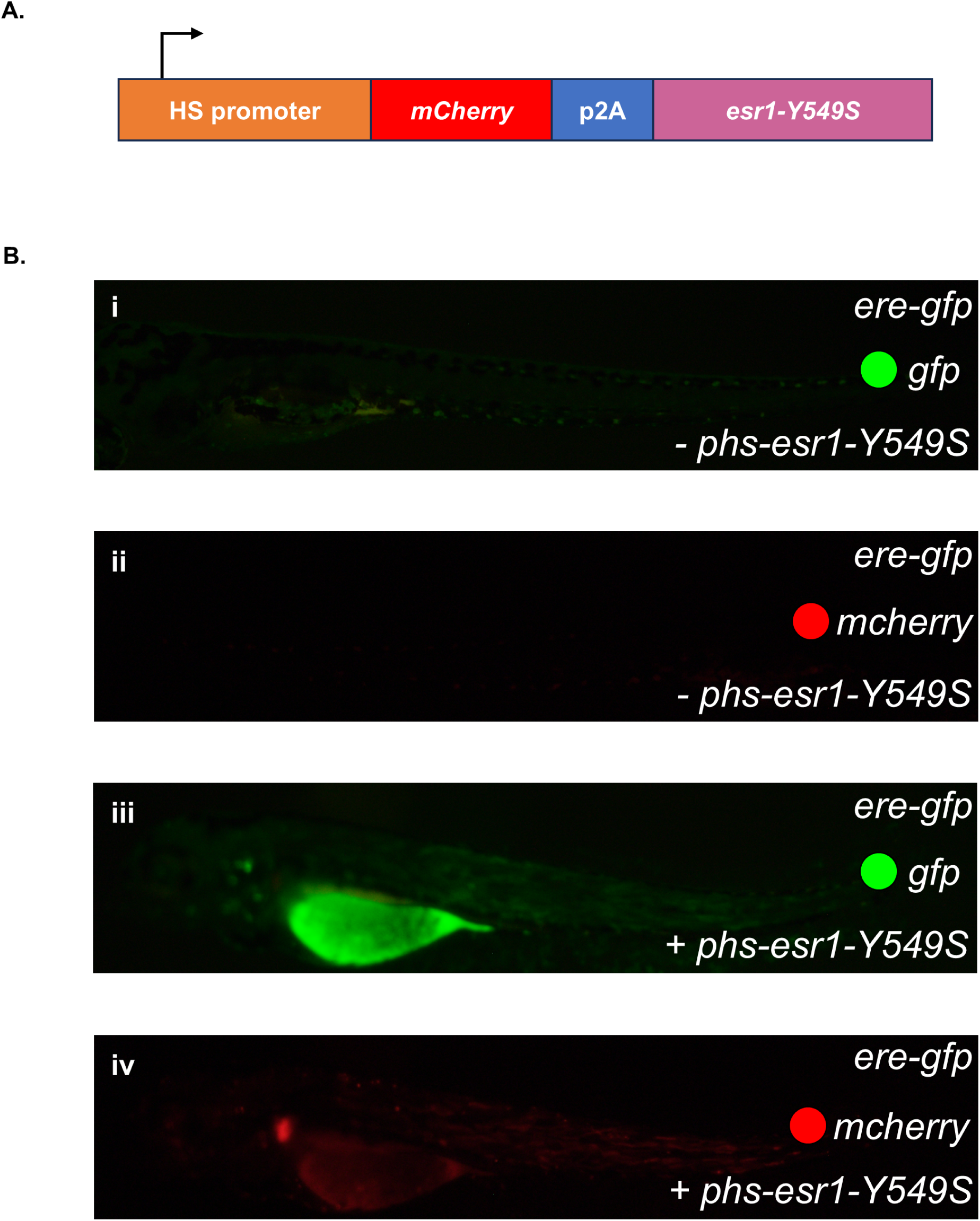
Constitutively active nuclear hormone estrogen receptor does not induce thrombosis. (A) p*hs*-*esr1*-Y549S plasmid with constitutively active *esr1* (Y549S). Expression was driven by the heat shock promoter (HS), along with *mcherry* followed by p2A and the mutant *esr1* cDNA. (B) p*hs*-*esr1*-Y549S was injected into *fgb-egfp;ere-gfp* transgenic embryos, treated at 35°C at 3 dpf, and evaluated at 5 dpf. GFP indicates activation of the estrogen response element (ERE), and mCherry indicates expression from p*hs*-*esr1*-Y549S. (i-ii) Uninjected controls show no ERE activity or mCherry expression. (iii-iv) Injected larvae show expression from p*hs*-*esr1*-Y549S by activation of ERE and subsequent GFP signal, as well as expression of mCherry. Despite evidence of constitutive *esr1* activity, no thrombosis was observed. All data were collected using the *fgb-egfp* transgenic background by an observer blinded to condition.

## Discussion

Thrombosis has been a well-known consequence of estrogen therapy for decades, yet we still know little about the underlying mechanisms. One of the major factors is the lack of an animal model. Blood coagulation is a complex process that relies on circulating cells and proteins in the context of an intact endothelial system, which is not easily modeled *in vitro*. Despite the aforementioned *in vitro* and *in vivo* murine studies, there remain many unanswered questions. We have developed a reliable and genetically tractable model using zebrafish, which has yielded some potential answers. Furthermore, this highly penetrant model that does not require weeks or months to develop a phenotype is a major advantage for future studies. Zebrafish treated with estrogens replicate key characteristics consistent with most human estrogen-induced thromboses, including typical development in the venous rather than arterial system, and lack of thrombosis in response to progesterone. Due to the transparency of zebrafish, we were also able to address a question that is essentially impossible to answer in mammals, i.e., the nature of the initiating event. In humans, thrombosis is often triggered by localized endothelial injury, e.g., trauma, surgery, and vascular access devices. Frequently, these thrombi subsequently spread through the vasculature. Since estrogen-induced thrombosis occurs weeks to months after initiation of therapy, it is difficult to determine if the process is similar. Through time-lapse imaging, we found that thrombosis initiates simultaneously along the venous endothelium. In our companion study, we also observed potential structural differences: estrogen-induced thrombi formed a speckled distribution, while in protein C (PC)-deficient lines (65) thrombi progressed via sprouting patterns with occasional dense aggregations (66). We hypothesize that estrogen has a direct effect on the endothelium, altering gene expression programs (67) that increase the risk of thrombosis.

Given what is known about estrogen signaling and our data, we believe that induction of thrombosis is preceded by alteration of transcription or cytoplasmic signaling in endothelial cells. We initially thought it would be easy to determine the first step through ablation of known receptors. However, through knockout and overexpression studies, we found that this effect is not mediated through canonical estrogen NHR or the known G-protein receptor. Despite these paradoxical findings, our data indicate that zebrafish thrombosis is a specific effect of estrogen, which does not occur with other steroid hormones and which is partially blocked by the estrogen antagonist fulvestrant. Finally, there is precedence for alternative targets or receptors. In elegant studies, Kelly and colleagues have clearly demonstrated the existence of a novel estrogen-responsive G-protein receptor, although the gene itself remains undiscovered (18). However, our data with STX suggest this is not our target. There is also evidence for other targets, including ion channels (10, 68). Finally, a recent zebrafish report also discovered that E2 promotes habituation learning through a novel target (19). It is tempting to speculate that estrogen-induced thrombosis may be mediated through one of these targets.

Surprisingly, estrogen-induced thrombosis persisted despite treatment with potent anticoagulants, as well as loss of common pathway procoagulant factors. We have previously shown that all of these medications and mutations completely block thrombosis in response to endothelial injury (34, 35, 38, 39). Similarly, in our companion drug-screening study, standard anticoagulants had little effect on estrogen-induced thrombosis yet effectively reduced spontaneous thrombosis driven by PC deficiency (66). These data indicate that estrogen-induced thrombi are not solely composed of fibrin secondary to thrombin-mediated cleavage of fibrinogen. However, the results of the a-chain knockout prove that the thrombi are indeed dependent on circulating fibrinogen. Thus, these results are not a fluorescence artifact, since it is known that ablation of *fga* results in absence of fibrinogen (36, 50), and our GFP tag is on the β-chain.

Interestingly, we found that estrogen-induced thrombosis depends on the TF paralogs, primarily *f3b*, and synergy when combined with *f3a*. This result contrasts with our previous study using a laser-endothelial injury model, where *f3b* drives arterial occlusion and *f3a* was primary in the venous system (52). However, that study also showed concordance with estrogen-induced thrombosis in that *f3b* was more effective for spontaneous venous thrombosis, which is known to be a consequence of antithrombin (*at3*) deficiency^34^, and was also synergistic with *f3a* (52). Laser-injury-induced thrombosis represents an acute hemostatic response, whereby damaged endothelial cells lead to vessel occlusion due to rapid recruitment of thrombocytes and formation of fibrin, in the arterial and venous systems, respectively. By contrast, estrogen-induced and spontaneous thromboses develop over hours to days, are independent of thrombocytes, and occur across and exclusively in the venous circulation, reflecting a chronic, hypercoagulable state rather than a site-specific injury. The data suggest the possibility of a resident *f3b* pool in the venous circulation that comes into play upon estrogen treatment without vascular injury. However, elevated levels of estrogen are known to affect inflammation (69, 70). Thus, an alternative is that TF is known to not be restricted to the endothelium, and monocytes, macrophages, and neutrophils also express TF under inflammatory conditions (71).

The next question we plan to address in future studies is the structure of these thrombi. Our data suggest several possibilities. The anticoagulants do inhibit thrombosis to a degree, so fibrin may be a partial component. Additionally, we found polyhedrocytes, which are known to form upon clot contraction (46). However, complete loss of the *f2* and *f10* genes should block all fibrin formation. Thus, there could be an alternative means of generating fibrin independent of thrombin, but this seems unlikely as there is no evidence for such a protease. A more likely hypothesis is that estrogen-induced thrombi are composed of crosslinked fibrinogen. It has previously been shown that non-factor XIII transglutaminases have this capability *ex vivo* (72). More recently, tissue transglutaminase-2 (TGM2) was shown to crosslink fibrin(ogen) in the setting of experimental liver fibrosis (73). Most intriguingly, injection of a Tgm2 fluorescent competitive substrate into zebrafish larvae yielded a labeling pattern (74) that bears a striking similarity to what we observe for estrogen-induced thrombosis. Tgm2 activity in zebrafish is primarily composed of two ohnologs (*tgm2a* and *b*), which were shown to be expressed during the larval period (1-5 dpf)(74).

We have found that estrogen-induced thrombosis in zebrafish shares many features consistent with human disease. Additionally, our findings reveal several unexpected aspects, including the involvement of a novel estrogen target/receptor, a connection with TF but not thrombin activity, and evidence of fibrin(ogen) crosslinking. Future studies could develop novel therapeutic targets as well as predict individuals at the highest risk for thrombosis.

## Methods

### Sex as a biological variable

All experiments were conducted using zebrafish embryos and larvae within the first 5 dpf. Since gonad differentiation in zebrafish begins after 10 dpf, sex is not a biological variable (75).

### Zebrafish

Zebrafish were maintained on a hybrid background of wild-type strains (ABxTL) to minimize strain-specific effects and enhance fecundity. Fish were examined at various developmental stages: embryo (0-2 days post-fertilization, dpf), larvae (3-29 dpf), juvenile (30-89 dpf), and adult (≥90 dpf) and housed at 28.5℃, with a 14-hour light/10-hour dark cycle. All procedures were approved by the University of Michigan Animal Care and Use Committee. Previously described transgenic and mutant lines include, *fgb-egfp* (labels fibrin(ogen) with green fluorescence)(76), prothrombin (*f2*)(39), factor X (*f10*)(35), *f3* (52), fibrinogen a-chain (*fga*)(36), *gper1*(60), *flk-mcherry* (red endothelium)(77), *ere-gfp* (estrogen response element reporter)(41), *moonshine* (*mon^tg234^*, absence of erythrocytes)(78), and *cd41-egfp* (79). Nearly all experiments were performed in the *fgb-egfp* background.

### Small molecule treatments

Chemicals were purchased from Sigma-Aldrich, dissolved in DMSO and then in E3 embryo buffer, followed by submersion of larvae in 2 ml E3 embryo buffer (Figure 1A). Final DMSO concentrations were always less than 2%. Larvae were treated at 3-5 dpf for 5-48 hours. Warfarin (50 µg/ml, Sigma-Aldrich) was initiated at 3 dpf and then followed until 5 dpf. Warfarin treatment was alternatively 10 µg/ml from 2 to 5 dpf, with mestranol (25 µM) added at 4 dpf to initiate thrombosis. Separate co-treatments of E2 (50 µM) with fulvestrant (50 µM), dabigatran etexilate (50 µM), or rivoraxaban (250 µM) were performed at 5 dpf for 5 hours. In addition, co-treatment of mestranol (2.5 µM) and fulvestrant (25 µM) was at 3 dpf for 48 hours.

### Imaging and quantification of thrombosis

After treatment, zebrafish larvae were anesthetized in tricaine and embedded in 0.8% low-melting point agarose on glass cover slips and visualized on an inverted microscope (Olympus IX71, 20x objective). For all experiments, the observer was blinded to treatment conditions and/or genotype using one of two methods. In the first approach, larvae-embedded slides were labeled and then masked with blank tags by the observer. A second investigator, blinded to the sample conditions, randomized and relabeled the slides in the absence of the observer. In the second method, images were obtained, saved with original filenames, then copied and renamed using randomly generated identifiers (sampling without replacement). In both methods, the key linking original and randomized identifiers was stored separately and accessed only after analysis was completed. In evaluating the thrombosis levels, a semi-quantitative grade was assigned based on the fluorescence intensity in the PCV. Grade 0 indicated no fluorescence, grade 1, sparse puncta, prominent accumulation in grade 2, and grade 3 exhibited dense, nearly complete coverage PCV (Figure 1B). Mutant larvae were recovered from agarose after evaluation and genotyped if needed. Confocal images were acquired by a Nikon A1 high sensitivity confocal microscope. Fluorescence intensity was measured with ImageJ to calculate CTCF as described(34). Concentration-response curves were generated based on both the ordinal thrombosis grading system and the CTCF quantification approach (66). Briefly, thrombosis values from each tested concentration were plotted on a log10-transformed concentration scale using a four-parameter nonlinear regression model (log[agonist] vs. response) in GraphPad Prism (v9.1.0), then normalized to a 0-100% scale.

### Measurement of plasma estrogen

After treatment, larvae were washed three times in E3 embryo buffer for 10 minutes, anesthetized, then individually bled in 5 µL of 4% sodium citrate diluted 1:6 in Tyrode’s buffer (80), with an estimated whole blood yield of 50 nL and thus 1:100 dilution. Ten larvae were pooled (50 µL), centrifuged at 500g for 10 minutes, and 30 µL supernatant was collected. This was diluted to 0.1 mL and mixed with 0.1-0.2 mL water and deuterium-labeled internal standards in 0.1 mL of 40% aqueous methanol. This mixture was applied to a supported liquid extraction cartridge (Isolute, Biotage) and analyzed by mass spectrometry as previously described(81), except that only the 1.50-2.65 min window from the first dimension chromatography was passed to the resolving column.

### Scanning electron microscopy (SEM)

After treatment with E2, larvae were immobilized using tricaine anesthesia and euthanized on ice. For SEM analysis, sagittal cuts were made in the larvae to open the PCV, rinsed in 50 mM sodium cacodylate buffer (pH 7.5) with 150 mM NaCl, and fixed in 2% glutaraldehyde. Samples were rinsed three times with the cacodylate buffer for 5 minutes and dehydrated in a series of increasing ethanol concentrations, immersed in hexamethyldisilazane, and dried overnight. A thin coating of gold-palladium was layered over the samples using a sputter coater (Polaron e5100, Quorum Technologies). Micrographs were captured using a Quanta 250 FEG scanning electron microscope (ThermoFisher Scientific).

### CRISPR-mediated genome editing

We used CRISPR/Cas9 and designed single guide RNAs (sgRNAs) to target upstream and downstream exons, thereby deleting nearly the entire locus of each *esr* gene (Table 1, Supplemental Figure 1) as previously described (52, 82). sgRNAs were complexed with Cas9 protein and injected into one-cell embryos. The *esr1* and *esr2a* genes are tightly linked, therefore *esr1* was targeted first, followed by *esr2a* once the former line had been established. Once the *esr1;esr2a* double mutants were produced, they were crossed to *esr2b* mutants to produce triple heterozygotes. *esr1;esr2a;esr2b* triple heterozygotes were incrossed to produce triple homozygous mutant larvae. Genotypes were confirmed using PCR primers (Table 2) and Sanger sequencing. sgRNAs were also designed to create a large deletion that spans the tandem *f7l* and *f7* loci (factors VII-like and VII, respectively), from exon 1 of *f7l* to exon 8 of *f7*, producing a double knockout.

**Table 1.**
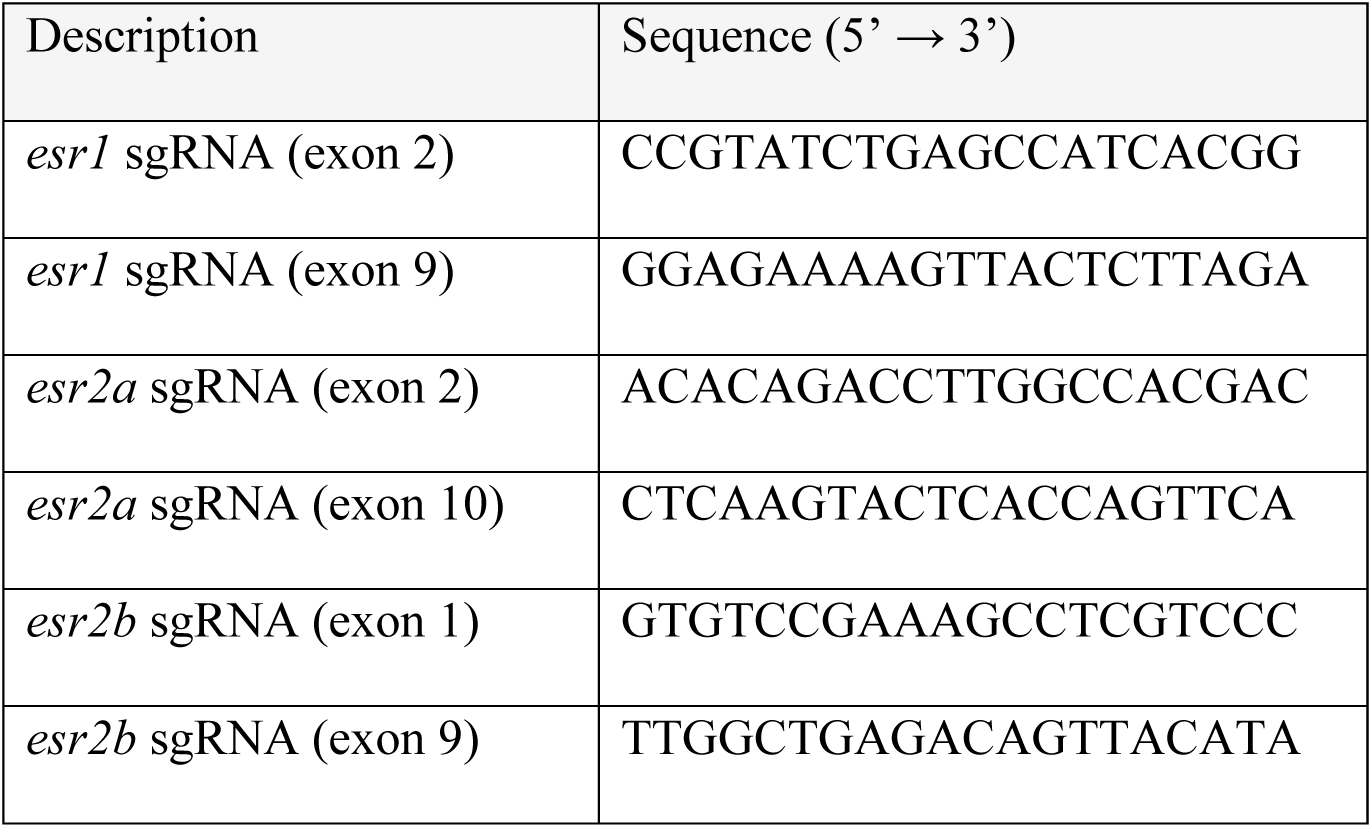
sgRNA sequences.

**Table 2.**
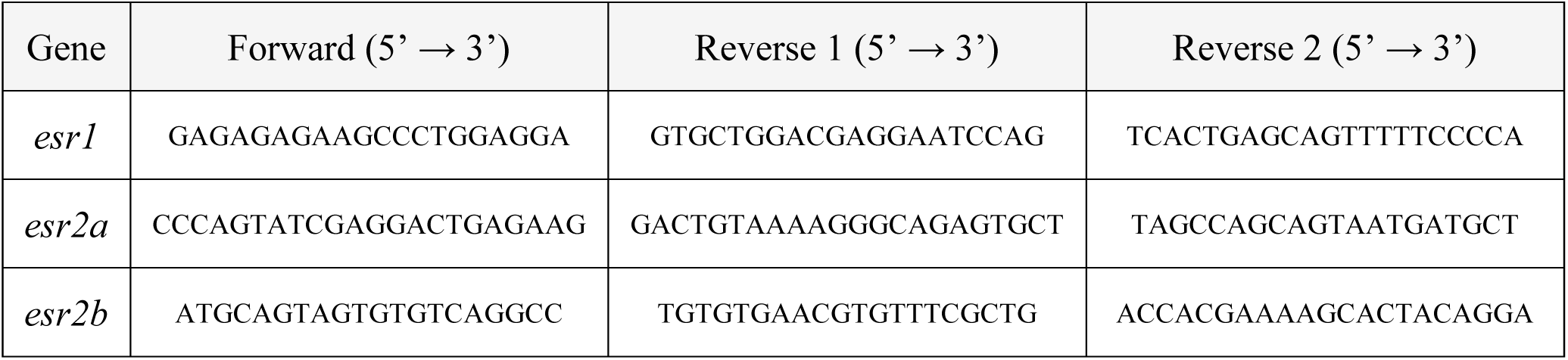
Primer sequences for genotyping.

### Transgene expression

The *esr1* overexpression plasmid (p*hs*-*esr1*-Y549S) was constructed by inserting the *esr1*-Y549S cDNA into a vector under control of the heat shock promoter(83) downstream of *mcherry* and a p2A sequence, all flanked by Tol2 repeats. 20-25 picograms plasmid was injected into *fgb-egfp* and *ere-gfp* one-cell stage embryos. At 3 dpf, larvae were incubated at 35°C and evaluated for thrombosis.

### Statistical analysis

Statistical analyses were performed using two-tailed Fisher exact, Student t-test and Spearman’s rank correlation test. Charts were generated using GraphPad Prism (v9.1.0, San Diego, CA, USA) and R (v4.3.3).

### Study approval

All experiments were conducted in accordance with guidelines approved by the Institutional Animal Care and Use Committee (IACUC) of the University of Michigan (Protocol Numbers: PRO00010679 & PRO00012352).

## Data availability

Data for all graphs are provided in the Supporting Data Values supplemental file.

## Authorship contributions

X.Y. and M.Y. designed and performed research, analyzed data, and wrote the manuscript. X.Y. and J.A.S. conceived of the project, and this was used to decide the order of co-authorship. Q.Y.Z., S.M.E., J.K.L., H.S. performed research and analyzed data. A.C.F., C.N., S.C. performed research, W.G., J.W.W., R.A. designed and supervised research and analyzed data, J.A.S. designed, performed, and supervised research, analyzed data, and wrote the manuscript. All authors reviewed the manuscript.

## Acknowledgements

The authors thank Martin Kelly for helpful discussions and providing STX compound, and Dan Gorelick for providing the *ere-gfp* line. This work was supported by National Institutes of Health grants R01 ES032255, R35 HL150784 (J.A.S.), R01 DK090311, R24 OD035402 (W.G.), and R01 HL148227, P01 HL146373, R01 HL148014, R01 HL159256 (J.W.W.), and a pilot grant from the University of Michigan Frankel Cardiovascular Center M-BRISC program. J.A.S. is the Henry and Mala Dorfman Family Professor of Pediatric Hematology/Oncology.

