## Supplementary figures and images for "Hormone-induced thrombosis is mediated through non-canonical fibrin(ogen) aggregation and a novel estrogen target in zebrafish"

### Supplemental Figure 1

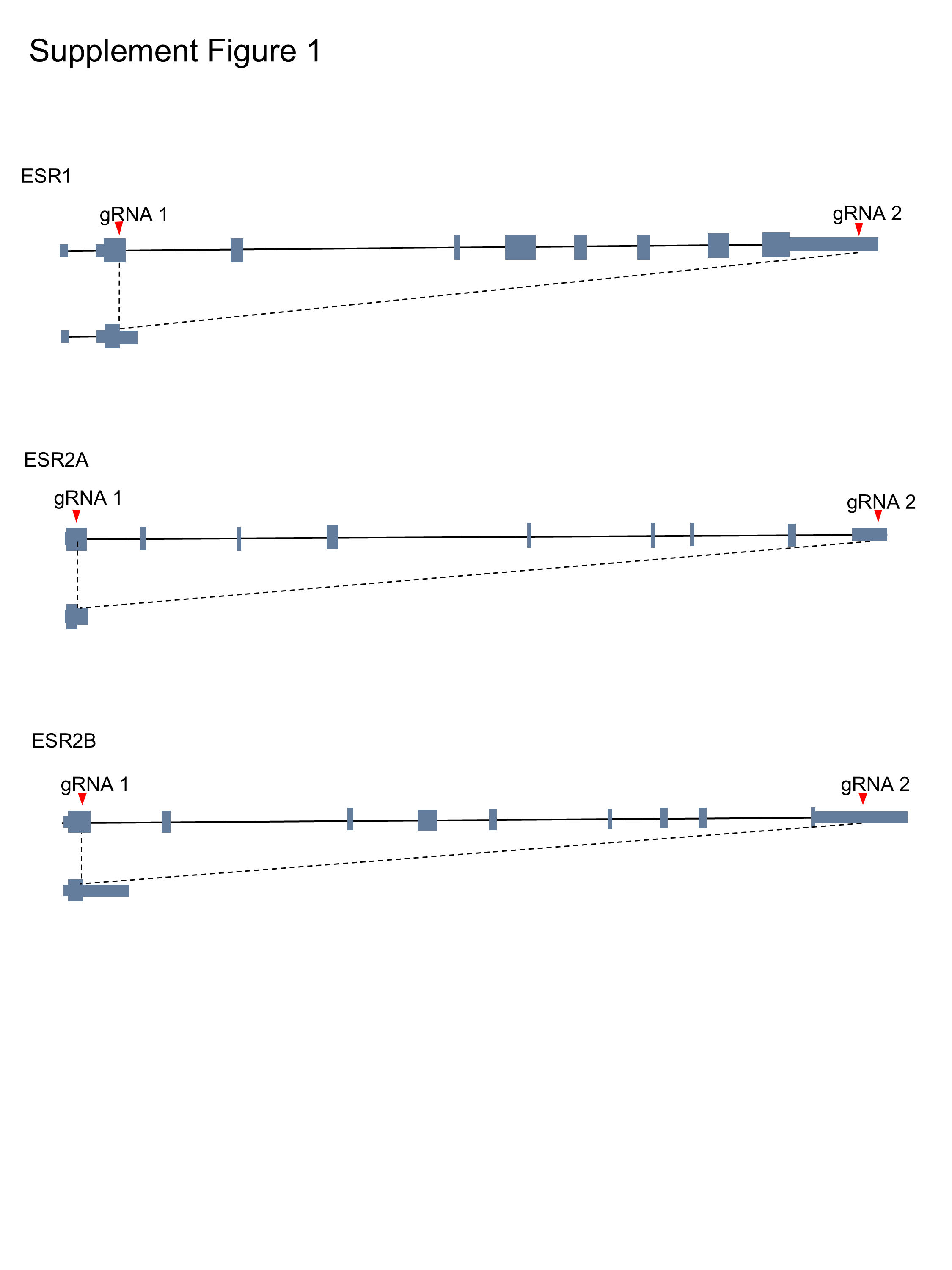
